## Supplementary material for "Maintenance of cell wall remodeling and vesicle production are connected in *Mycobacterium tuberculosis*": FigS1-13

**A**

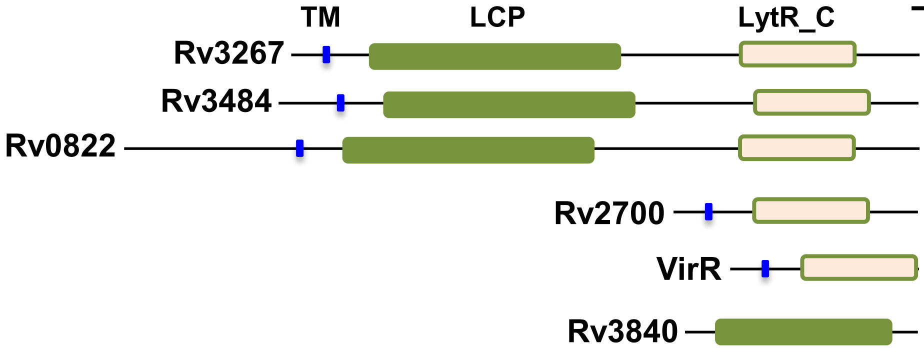


| | Growth defect | Release material | Attenuated Virulence |
|---|---|---|---|
| Rv3267 | Yes | No | No |
| Rv3484 | No | No | Yes |
| Rv0822 | No | No | No |
| Rv2700 | Yes | ? | Yes |
| VirR | -/+ | +++ | Yes |
| Rv3840 | ? | ? | ? |

**B**

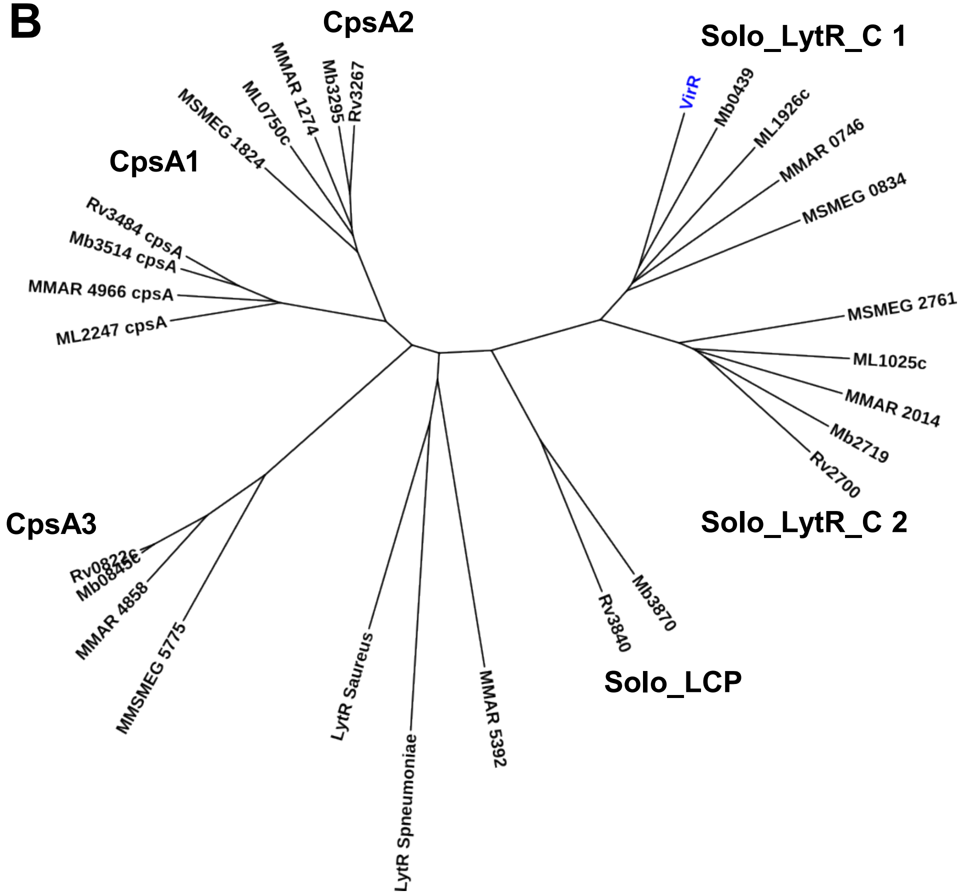


Fig S1

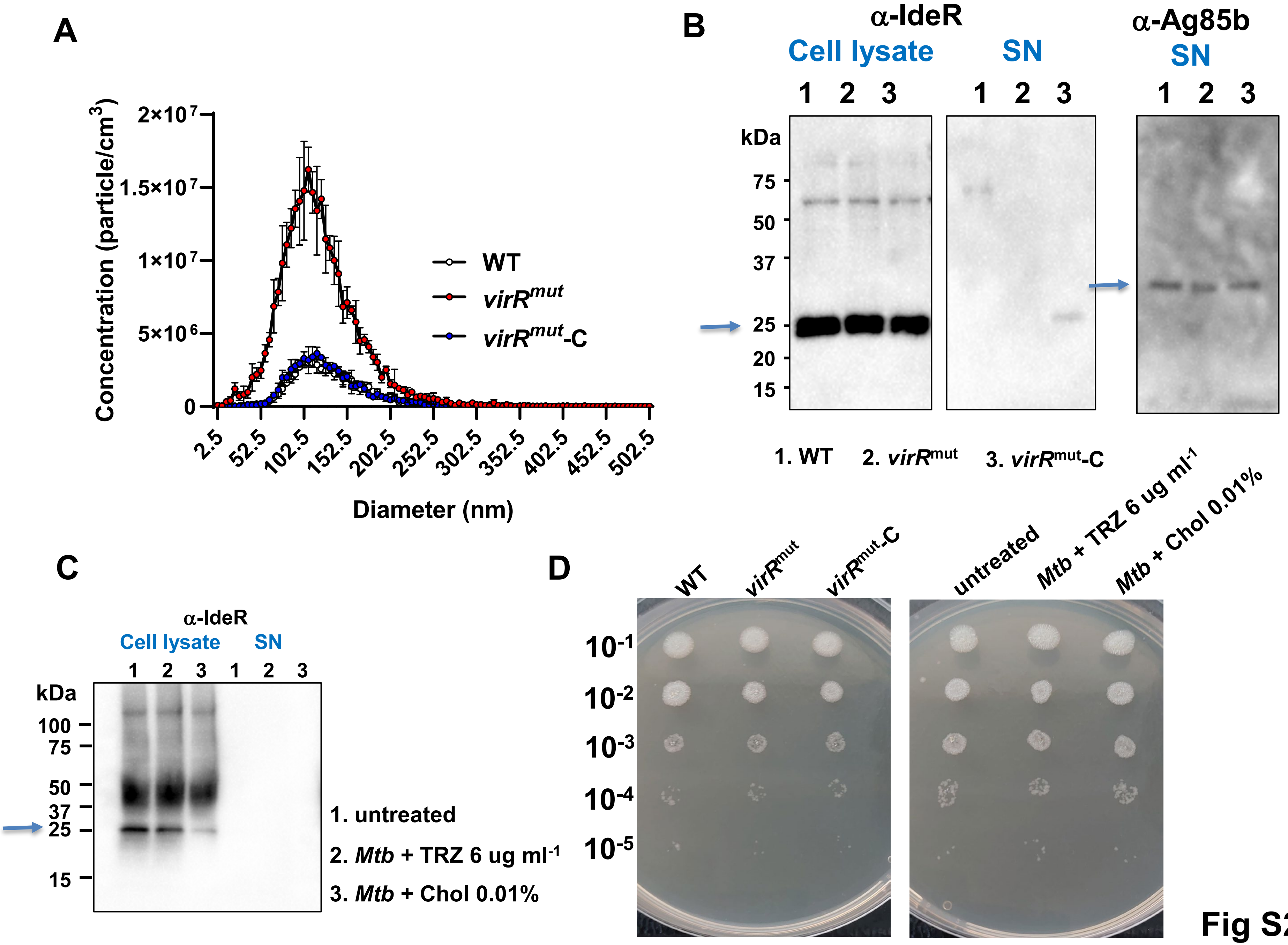
A
Concentration (particle/cm3)
2×10^7
1.5×10^7
1×10^7
5×10^6
0
2.5
52.5
102.5
152.5
202.5
252.5
302.5
352.5
402.5
452.5
502.5
Diameter (nm)
WT
virRmut
virRmut-C
B
α-IdeR
Cell lysate
SN
α-Ag85b
SN
1 2 3
1 2 3
1 2 3
kDa
75
50
37
25
20
15
1. WT
2. virRmut
3. virRmut-C
C
α-IdeR
Cell lysate
SN
1 2 3 1 2 3
kDa
100
75
50
37
25
15
1. untreated
2. Mtb + TRZ 6 ug ml-1
3. Mtb + Chol 0.01%
D
WT
virRmut
virRmut-C
untreated
Mtb + TRZ 6 ug ml-1
Mtb + Chol 0.01%
10-1
10-2
10-3
10-4
10-5

Fig S2

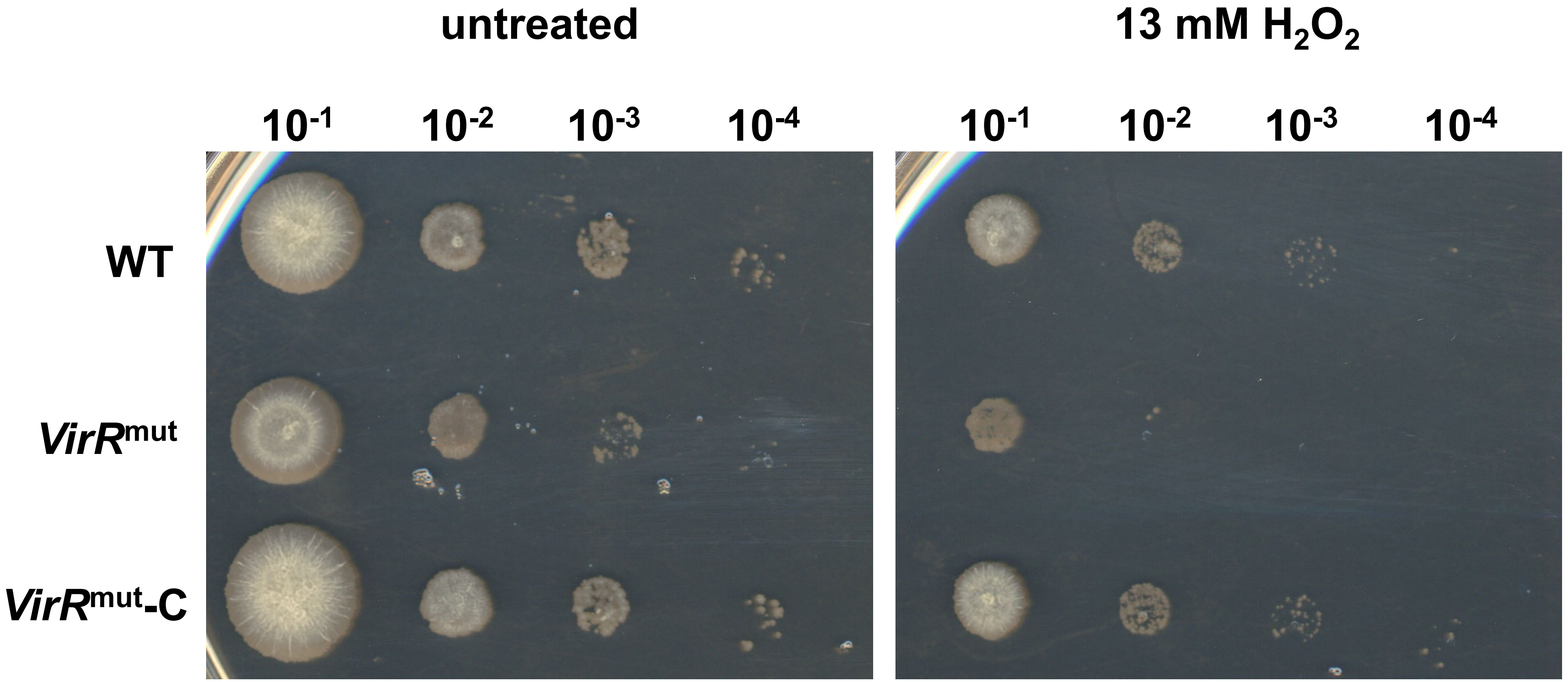
untreated
13 mM H2O2
10-1
10-2
10-3
10-4
10-1
10-2
10-3
10-4
WT
VirRmut
VirRmut-C

Fig S3

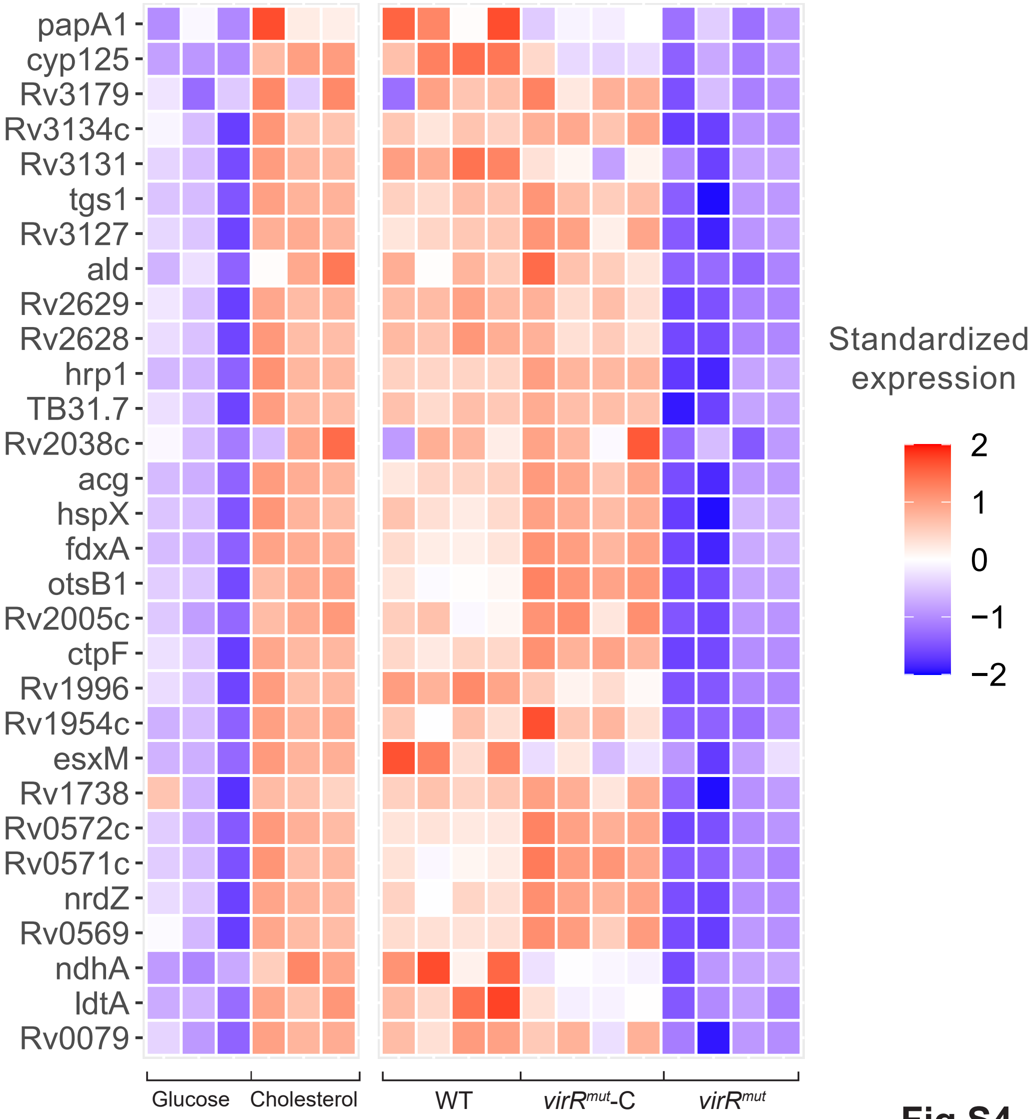
papA1
cyp125
Rv3179
Rv3134c
Rv3131
tgs1
Rv3127
ald
Rv2629
Rv2628
hrp1
TB31.7
Rv2038c
acg
hspX
fdxA
otsB1
Rv2005c
ctpF
Rv1996
Rv1954c
esxM
Rv1738
Rv0572c
Rv0571c
nrdZ
Rv0569
ndhA
ldtA
Rv0079
Glucose
Cholesterol
WT
virR^mut-C
virR^mut
Standardized expression
2
1
0
−1
−2

**Fig S4**

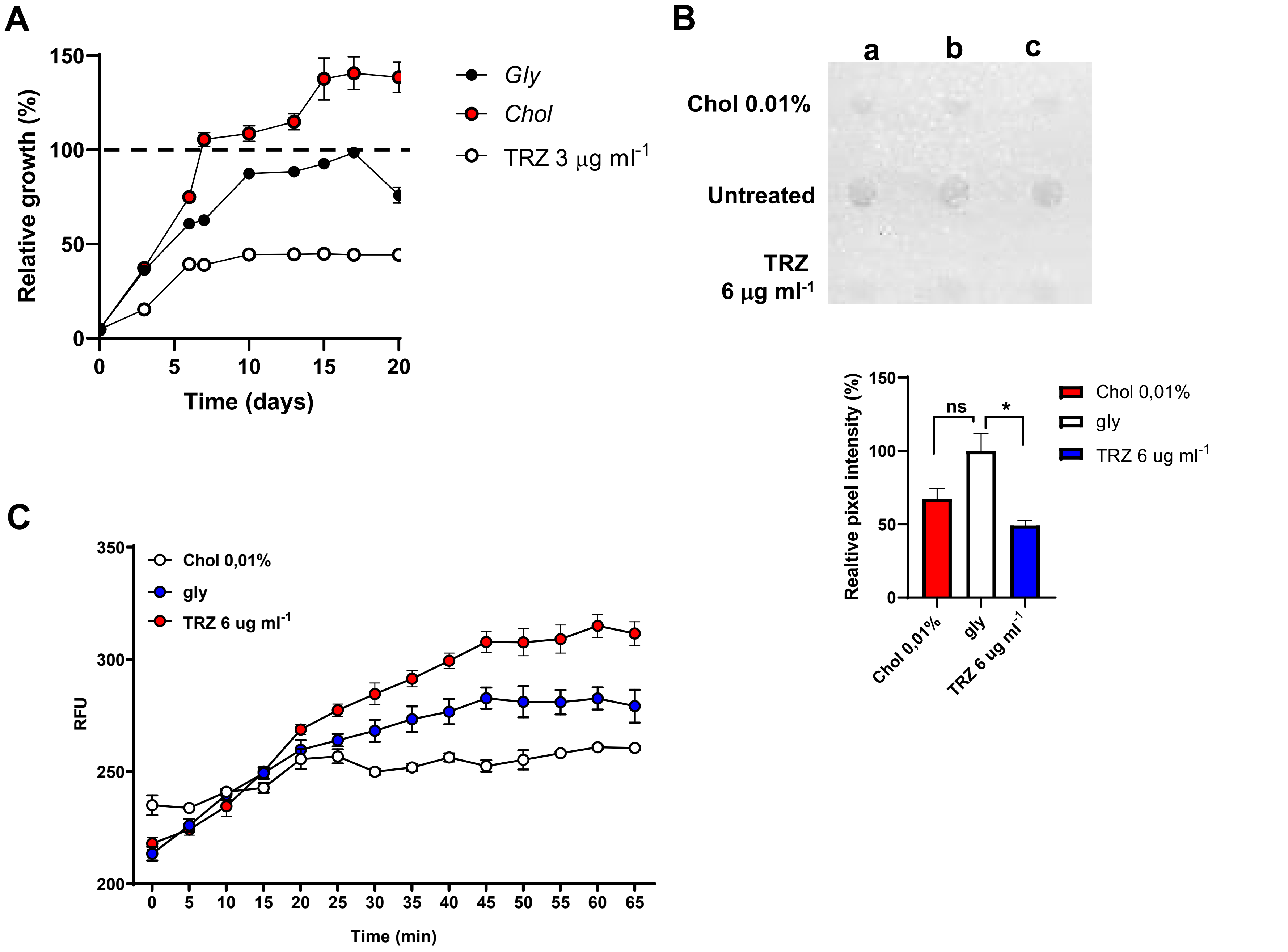
A
Relative growth (%)
150
100
50
0
0
5
10
15
20
Time (days)
Gly
Chol
TRZ 3 μg ml-1
B
a
b
c
Chol 0.01%
Untreated
TRZ
6 μg ml-1
Realtive pixel intensity (%)
ns
*
Chol 0,01%
gly
TRZ 6 ug ml-1
C
350
300
250
200
RFU
Chol 0,01%
gly
TRZ 6 ug ml-1
0
5
10
15
20
25
30
35
40
45
50
55
60
65
Time (min)

Fig S5

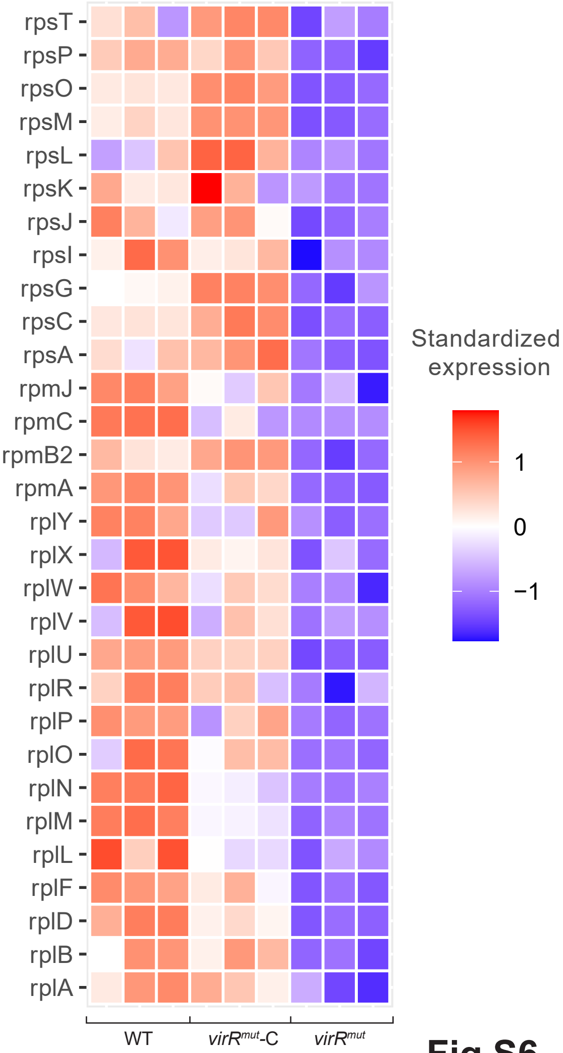
rpsT
rpsP
rpsO
rpsM
rpsL
rpsK
rpsJ
rpsI
rpsG
rpsC
rpsA
rpmJ
rpmC
rpmB2
rpmA
rplY
rplX
rplW
rplV
rplU
rplR
rplP
rplO
rplN
rplM
rplL
rplF
rplD
rplB
rplA
WT
virRmut-C
virRmut
Standardized expression
1
0
−1

**Fig S6**

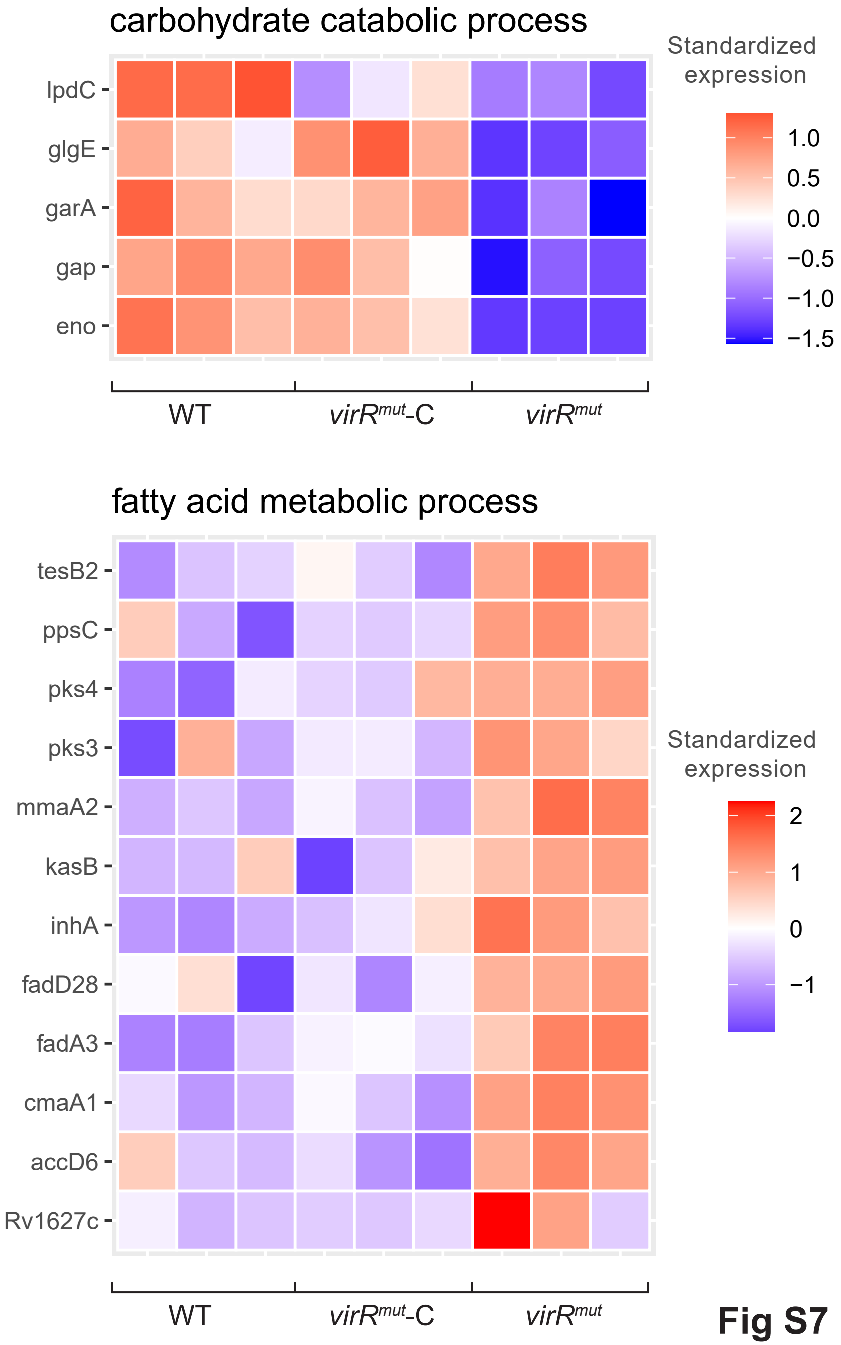
carbohydrate catabolic process
Standardized expression
1.0
0.5
0.0
−0.5
−1.0
−1.5
lpdC
glgE
garA
gap
eno
WT
*virR^mut^-C*
*virR^mut^*
fatty acid metabolic process
Standardized expression
2
1
0
−1
tesB2
ppsC
pks4
pks3
mmaA2
kasB
inhA
fadD28
fadA3
cmaA1
accD6
Rv1627c
WT
*virR^mut^-C*
*virR^mut^*

**Fig S7**

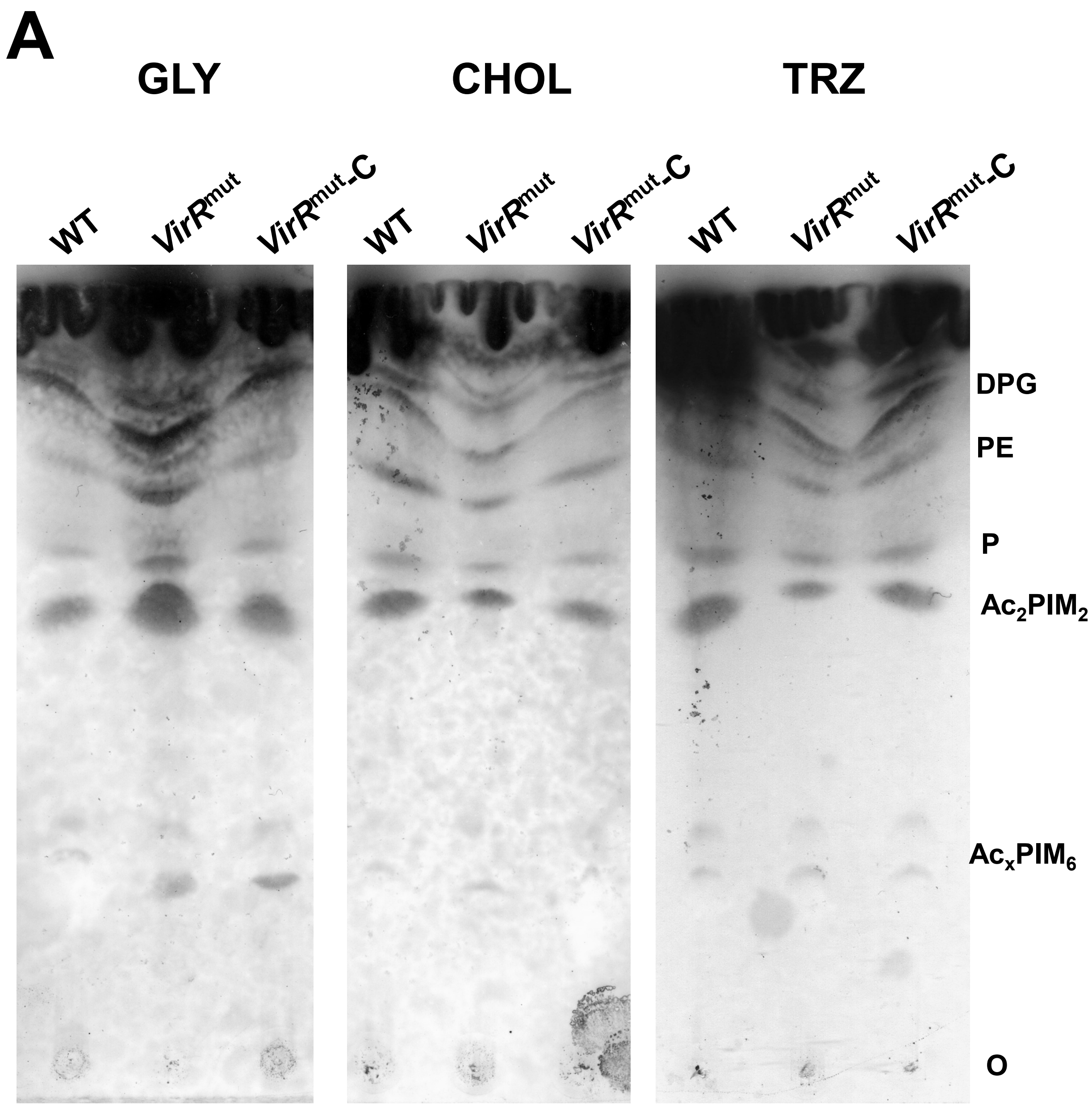
A
GLY
CHOL
TRZ
WT
VirR^mut
VirR^mut-C
WT
VirR^mut
VirR^mut-C
WT
VirR^mut
VirR^mut-C
DPG
PE
P
Ac2PIM2
AcxPIM6
O

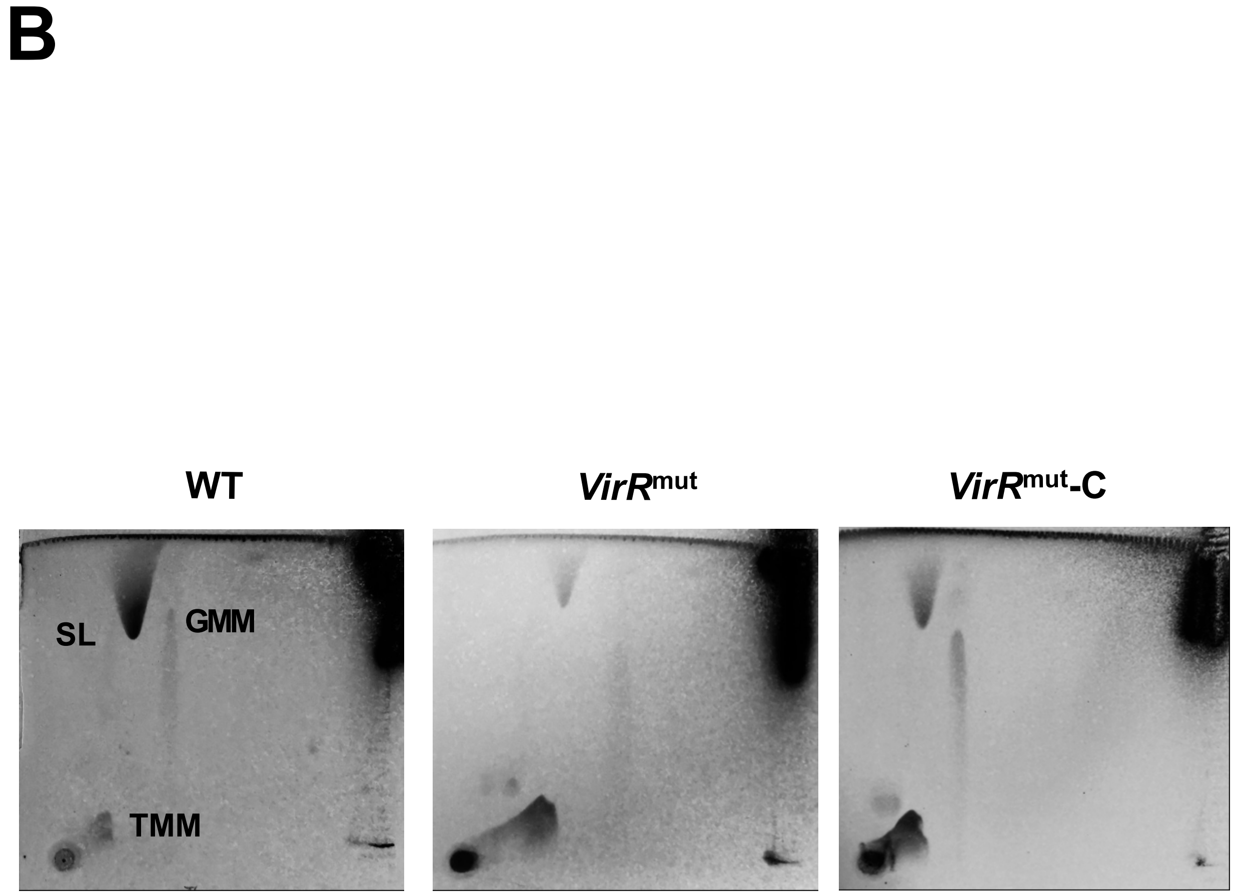
B
WT
VirR^mut
VirR^mut-C
SL
GMM
TMM

FigS8

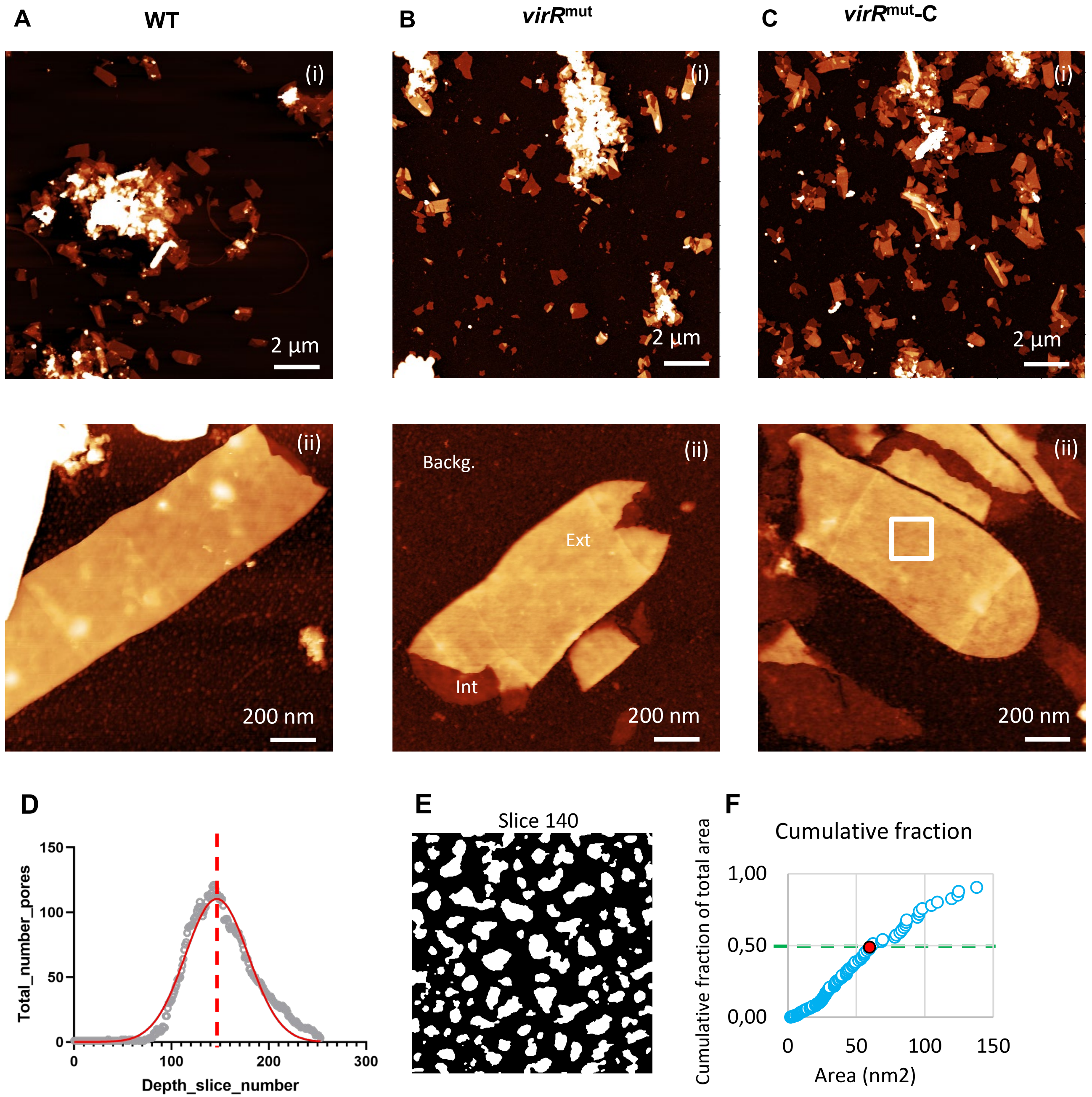
A
WT
B
virR^mut
C
virR^mut-C
(i)
2 µm
(ii)
200 nm
Backg.
Ext
Int
D
Total_number_pores
Depth_slice_number
0
50
100
150
200
300
E
Slice 140
F
Cumulative fraction
Cumulative fraction of total area
1,00
0,50
0,00
Area (nm2)

Fig S9

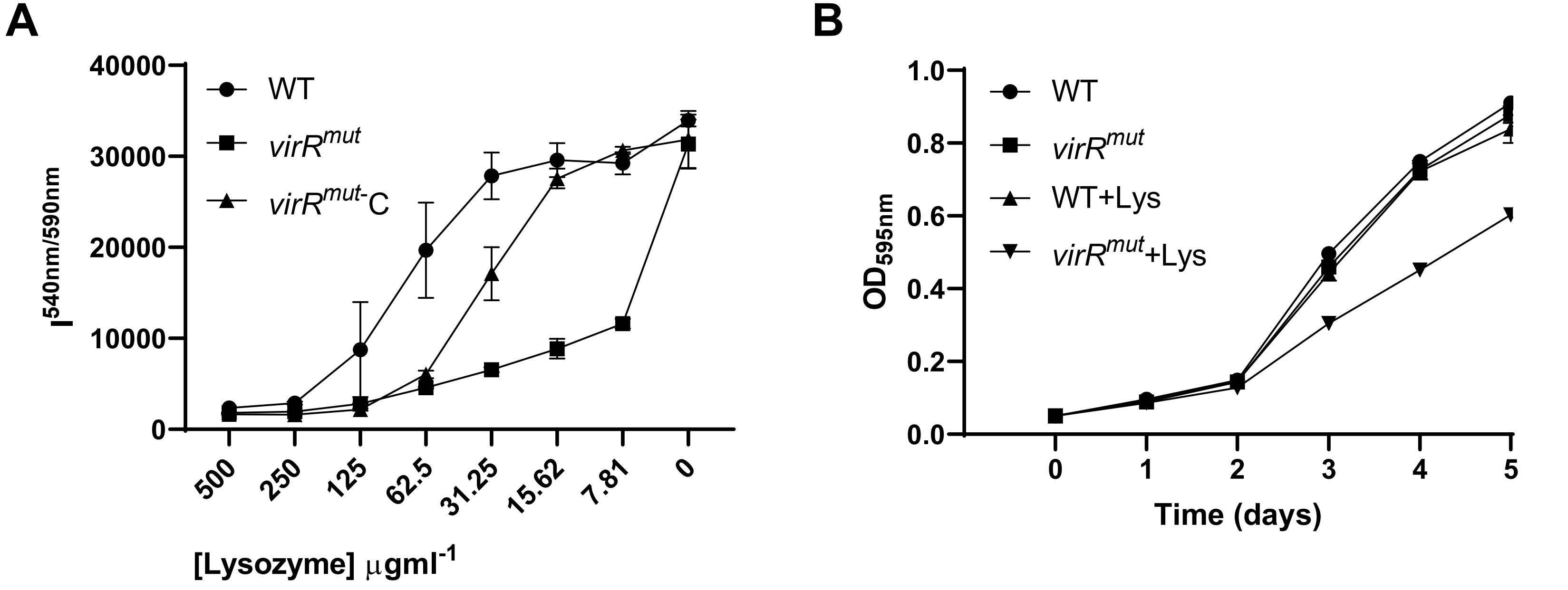
A
B
WT
virR^mut
virR^mut-C
WT+Lys
virR^mut+Lys
I^540nm/590nm
40000
30000
20000
10000
0
500
250
125
62.5
31.25
15.62
7.81
0
[Lysozyme] μgml^-1
OD595nm
1.0
0.8
0.6
0.4
0.2
0.0
0
1
2
3
4
5
Time (days)

Fig S10

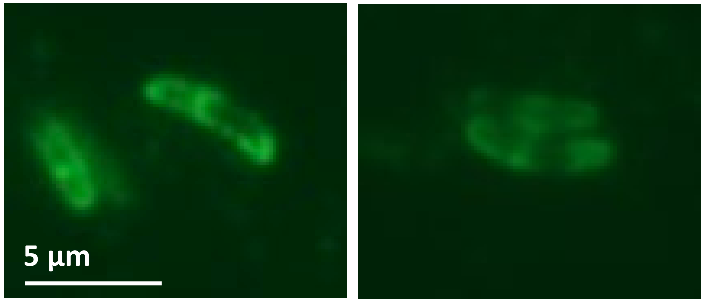


Fig S11

A

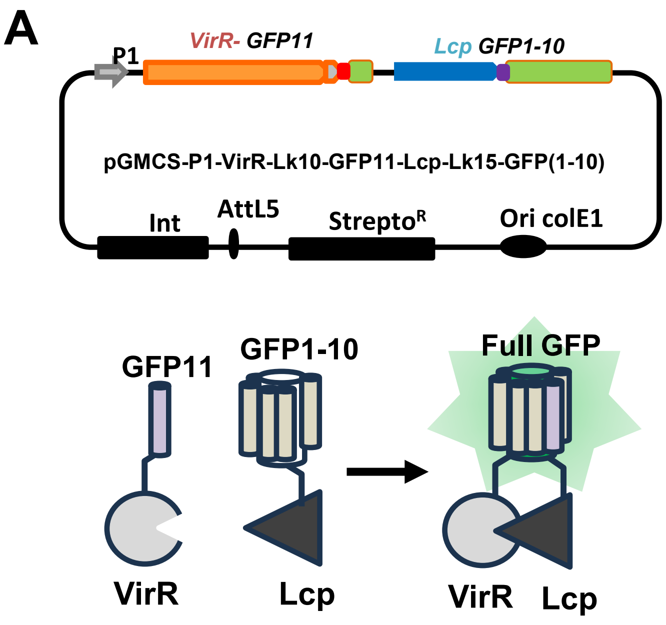
P1
VirR- GFP11
Lcp GFP1-10
pGMCS-P1-VirR-Lk10-GFP11-Lcp-Lk15-GFP(1-10)
Int
AttL5
StreptoR
Ori colE1
GFP11
GFP1-10
Full GFP
VirR
Lcp
VirR
Lcp

B

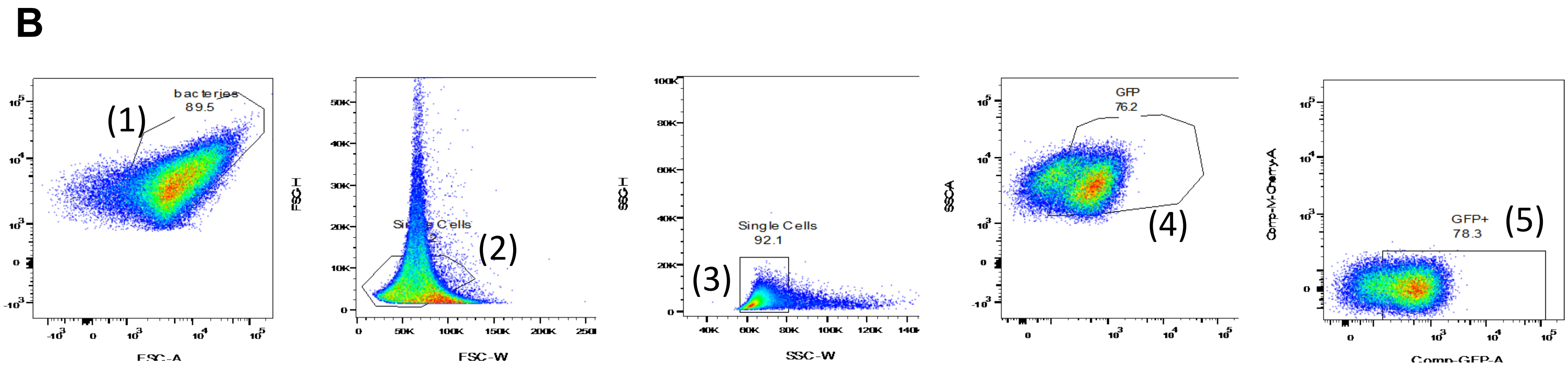
(1)
bacteries
89.5
FSC-A
(2)
Single Cells
FSC-W
FSC-H
(3)
Single Cells
92.1
SSC-W
SSC-H
GFP
76.2
(4)
SSC-A
GFP+
78.3
(5)
Comp-M-Cherry-A
Comp-GFP-A

Fig S12

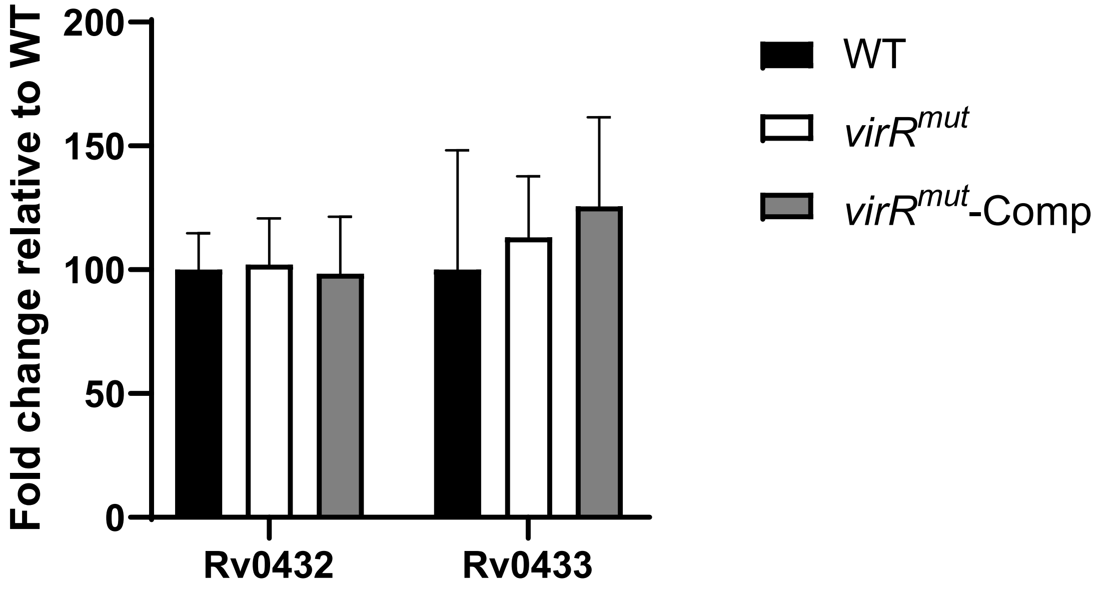
Fold change relative to WT
200
150
100
50
0
Rv0432
Rv0433
WT
virR^mut
virR^mut-Comp

**Fig S13**
